# Transcriptional Profiling and Co-expression Integration for the Filtering of Relevant Bacterial sRNA–mRNA Interactions: Application to *Staphylococcus aureus* Biofilm

**DOI:** 10.1101/2025.11.26.690695

**Authors:** Carolina Albuquerque Massena Ribeiro, Guadalupe del Rosario Quispe Saji, Maiana de Oliveira Cerqueira e Costa, Alice Slotfeldt Viana, Mariana Fernandes Carvalho, Agnes Marie Sá Figueiredo, Edgardo Galán-Vásquez, J. Eduardo Martinez-Hernandez, Marisa Fabiana Nicolas

## Abstract

Small regulatory RNAs (sRNAs) are fast-acting non-coding RNAs (ncRNAs), stress-responsive regulators that fine-tune bacterial gene expression, shaping virulence, antimicrobial resistance, metabolism, and biofilm development. At the post-transcriptional level, sRNAs pair with target mRNAs to block or enhance translation, remodel secondary structures, adjust transcript stability, and act as molecular sponges for other sRNAs. *Staphylococcus aureus*, a leading cause of hospital-acquired infections, relies on a multiple-layered regulatory network, including post-transcriptional mechanisms, to transition between planktonic and biofilm lifestyles. Here, we expand the cross-lineage sRNA repertoire of *S. aureus* by integrating newly generated RNA-seq data from the Brazilian ST239 strain Bmb9393 with public datasets from five USA-lineage strains previously uncharacterized for sRNAs. Using sequence homology and covariance models, we predicted and annotated candidate sRNA loci across all analyzed genomes, quantified their expression under planktonic and biofilm conditions, and assigned genomic context. Integration of differential-expression (DE) profiles with weighted gene co-expression network analysis (WGCNA) identified sRNAs associated with biofilm and virulence, in modules that include well-known regulators (*sarA, mgrA*, RNAIII) and the adhesin *clfA*. To prioritize functional target interactions, we combined DEG concordance, network features, and interaction-energy thresholds, depleting millions of initial predictions to thousands of high-confidence sRNA–mRNA pairs. Our integrative bioinformatics framework provides additional insights into sRNA-mediated regulation in *S. aureus*, highlighting biofilm- and resistance-linked candidates, and yields a ranked, reusable set of sRNA–mRNA interactions to guide hypothesis-driven experiments across diverse genetic backgrounds.

## Introduction

Antimicrobial resistance remains a significant global public health concern, with methicillin-resistant *S. aureus* (MRSA) representing one of the most critical threats. In Latin America, MRSA accounts for a substantial share of *S. aureus* infections. Regional estimates indicate that at least one quarter of isolates are methicillin-resistant. In Brazil, national surveillance by ANVISA has consistently documented a high burden of MRSA in adult intensive care units (ICUs) and in catheter-related bloodstream infections, with recent reports showing more than half of *S. aureus* bloodstream isolates from adult ICUs. This burden is amplified in ICUs, where the widespread use of invasive devices such as central venous catheters predisposes patients to biofilm-associated infections, highlighting the urgent need for strategies that effectively address biofilm-associated infection and resistance (Seas *et al*., 2018; *Boletins e relatórios — Agência Nacional de Vigilância Sanitária - Anvisa*, 2025).

Historically, antibacterial drug development has focused on protein-coding targets in essential pathways (e.g., cell wall, replication, metabolism), and this paradigm remains the backbone of therapy (Nicolás *et al*., 2020). However, repeated antibiotic exposure accelerates selection for resistance and shortens the clinical lifespan of antimicrobials (Belete, 2019; Serral *et al*., 2021). In light of rising bacterial resistance, therapeutic discovery should diversify the target portfolio, incorporating underexplored coding genes alongside ncRNA-based regulatory pathways. Research on ncRNAs, especially sRNAs, offers promising avenues for combating multidrug-resistant bacteria, such as MRSA and Enterobacteriaceae. These sRNAs are rapid-acting, stress-responsive post-transcriptional regulators that modulate microbial metabolism and other cellular processes. Depending on target and context, they can block translation by via base-pairing to occlude the Shine– Dalgarno (SD) sequence; boost translation by exposing the SD; either promote or prevent target mRNA degradation; and act as molecular sponges by sequestering other sRNAs or RNA-binding proteins (Bouvier *et al*., 2008; Desnoyers, Bouchard and Massé, 2013; Carrier, Lalaouna and Massé, 2018; Dutta and Srivastava, 2018; Leistra, Curtis and Contreras, 2019), (Ribeiro *et al*., 2024).

Therapeutic strategies that harness or target sRNAs to reprogram bacterial gene expression constitute a novel, emerging anti-infective approach. A proof-of-concept has been demonstrated with small molecules that bind structured RNAs and modulate function. For example, ribocil, a phenotypically discovered synthetic mimic of flavin mononucleotide, selectively engages the riboflavin riboswitch to repress *ribB* expression and inhibit bacterial growth, underscoring that the ncRNA elements may be broadly druggable (Howe *et al*., 2015). This approach moves beyond conventional paradigms that focus solely on druggable protein targets, expanding the repertoire of potential therapeutic targets with novel modes of action (MOAs) (Dersch *et al*., 2017; Matsui and Corey, 2017; Parmeciano Di Noto, Molina, and Quiroga, 2019; Wang *et al*., 2022). The versatility of sRNAs, combined with their ability to coordinate the regulation of multiple targets, positions them as a powerful tool in combating antimicrobial resistance (Howden *et al*., 2013; Dersch *et al*., 2017; Lalaouna *et al*., 2019; Mediati *et al*., 2021).

High-throughput RNA sequencing has revolutionized bacterial transcriptomics, enabling the systematic and cost-effective discovery of regulatory ncRNAs. Public genome and transcriptome resources, coupled with modern bioinformatics, now support detailed characterization of sRNAs and their interactions. Integrating public datasets with *in silico* approaches provides valuable insights into leveraging these molecules as therapeutic targets. Although these datasets are available, many remain untapped for sRNA-focused analysis, representing a valuable but largely unexplored reservoir of knowledge (Kim, 2017; Batool, Ahmad, and Choi, 2019; Liao *et al*., 2022; X. Zhang *et al*., 2022). In this context, prediction *in silico* of mRNA targets of sRNA is essential for advancing and scaling this research area. Methods that integrate thermodynamic modeling, comparative genomics, and machine learning can prioritize candidate pairs of sRNA-mRNA across entire genomes. However, computational predictions of their targets have intrinsic limitations: context-dependent RNA dynamics, RNA-binding proteins, and modifications can lead to false positives and negatives. Therefore, pipelines must integrate multiple layers of evidence and prioritize candidates for empirical validation (e.g., in vitro/in vivo) to establish function (Li, Ying, and Cha, 2012; Georg *et al*., 2020).

To address these challenges, we combined newly generated RNA-seq data for the Brazilian ST239 strain Bmb9393 of *S. aureus* with reanalyzed public datasets from five USA-lineage strains to prioritize robust sRNA–mRNA interactions. We mapped loci, quantified expression under planktonic and biofilm-promoting conditions, and integrated DEG with WGCNA to recover regulatory modules. We applied constraints based on interaction energy thresholds, DEG, and co-expression to filter mRNA-target predictions of the identified sRNAs. Accordingly, we deliver a prioritized, high-confidence dataset of sRNA–mRNA interactions that reveal pathways associated with biofilm formation, stress responses, and metabolism, providing tractable hypotheses for targeted testing across diverse genetic backgrounds and conditions.

## Results and Discussion

### Identification of sRNAs

The first significant outcome of this study is a genome-wide annotation of non-coding sRNAs in the *S. aureus* Bmb9393 genome, achieved here for the first time. Additionally, an in-house Python script was developed to categorize systematically predicted sRNA loci based on their genomic locations. The script was applied to five additional strains, providing a foundational framework for subsequent analyses of sRNA expression and target mRNA prediction (see Supplementary Tables S1-S5). Since a well-annotated reference genome for NRS385 USA500 is not yet available, we used the LAC USA300 reference genome, as it represents a closely related available genome, in agreement with previous reports (Li *et al*., 2009; Glaser *et al*., 2016; Tomlinson, Malof, and Shaw, 2021). A summary of the number of sRNAs identified in each category is presented in Table 1.

**Table 1:** Genetic Location of sRNA Classes Across Strains.

| Location | Bmb9393 | N315 | MRSA252 | LAC | MW2 |
| --- | --- | --- | --- | --- | --- |
| Antisense | 197 | 197 | 206 | 195 | 214 |
| Intragenic | 58 | 53 | 56 | 54 | 57 |
| 5' UTR | 63 | 56 | 50 | 51 | 55 |
| 3' UTR | 65 | 60 | 69 | 51 | 69 |
| Intergenic | 303 | 179 | 238 | 173 | 242 |

We next analyzed RNA-seq libraries (coding sequences and non-coding sequence loci) from the Bmb9393 strain together with previously published datasets of USA strains. To identify orthologous genes across all strains, we implemented the homology-based workflow using ProteinOrtho to infer shared and strain-specific gene sets and transfer functional annotations. Regarding coding sequences (CDSs) in *S. aureus* Bmb9393, we retrieved 2,934, and the compared strains contained 2,706 (MW2), 2,736 (N315), 2,802 (MRSA252), and 2,857 (LAC/NRS385) CDSs (see Supplementary Tables S6-S11). ProteinOrtho identified 2,330 orthologous groups shared by all strains, with Bmb9393 exhibiting the largest number of unique genes (n = 212). Ortholog assignments can be cross-checked in Supplementary Tables S6–S11, where we list, for each gene ID row, the corresponding locus tags from all strains whenever orthology is established. Overall, 80.42% of gene groups were conserved across strains belonging to ST239 and USA-lineage backgrounds.

In parallel, our RNA-seq analysis detected 291 sRNAs common to all strains, with sequence identity typically in the 70–80% range (Figure S1). To define the presence of an sRNA in our data, we applied a threshold requiring a mean of at least 10 reads across replicates, normalized by TMM and RPKM methods. This threshold was chosen based on the principle that low read counts are often associated with higher variability and reduced confidence in gene expression measurements, particularly for sRNAs. The number of sRNAs detected varied across strains, with Bmb9393 showing the highest count (512), followed by N315 (443), LAC (437), MW2 (433), NRS385 (406), and MRSA252 (404) (Figure S1).

### DEG results from paired-end RNA-Seq: coding genes and sRNAs

Our analysis of sRNA differential expression between biofilm and planktonic lifestyles highlights several state-specific sRNAs, consistent with context-dependent regulatory functions. We found 70 upregulated sRNAs DEGs common to all five strains, but this number decreases to 10 when Bmb9393 is included (Figure S2). Notably, we identified several that correlate strongly with biofilm growth upon investigating these two sets of sRNAs. Similarly, the set of downregulated sRNAs in biofilm includes 39 shared among the five strains excluding Bmb9393, but this number drops to just 4 when Bmb9393 is included in the analysis (Figure S2).

The heatmap in Figure 1 shows the 10 sRNAs consistently up- or downregulated across all strains. Specifically, ten sRNAs were consistently upregulated, including rsaH, sRNA160, sRNA4, Teg13, sprG3, sprF3, artR, Sau6873, Sau92, and Teg38as, while four sRNAs were consistently downregulated, namely Tegg112, RsaX05, sRNA47, and Sau14. This consistent expression suggests that specific sRNAs may play conserved regulatory roles in biofilm and planktonic conditions across the *S. aureus* lineages studied.

**Figure 1.**
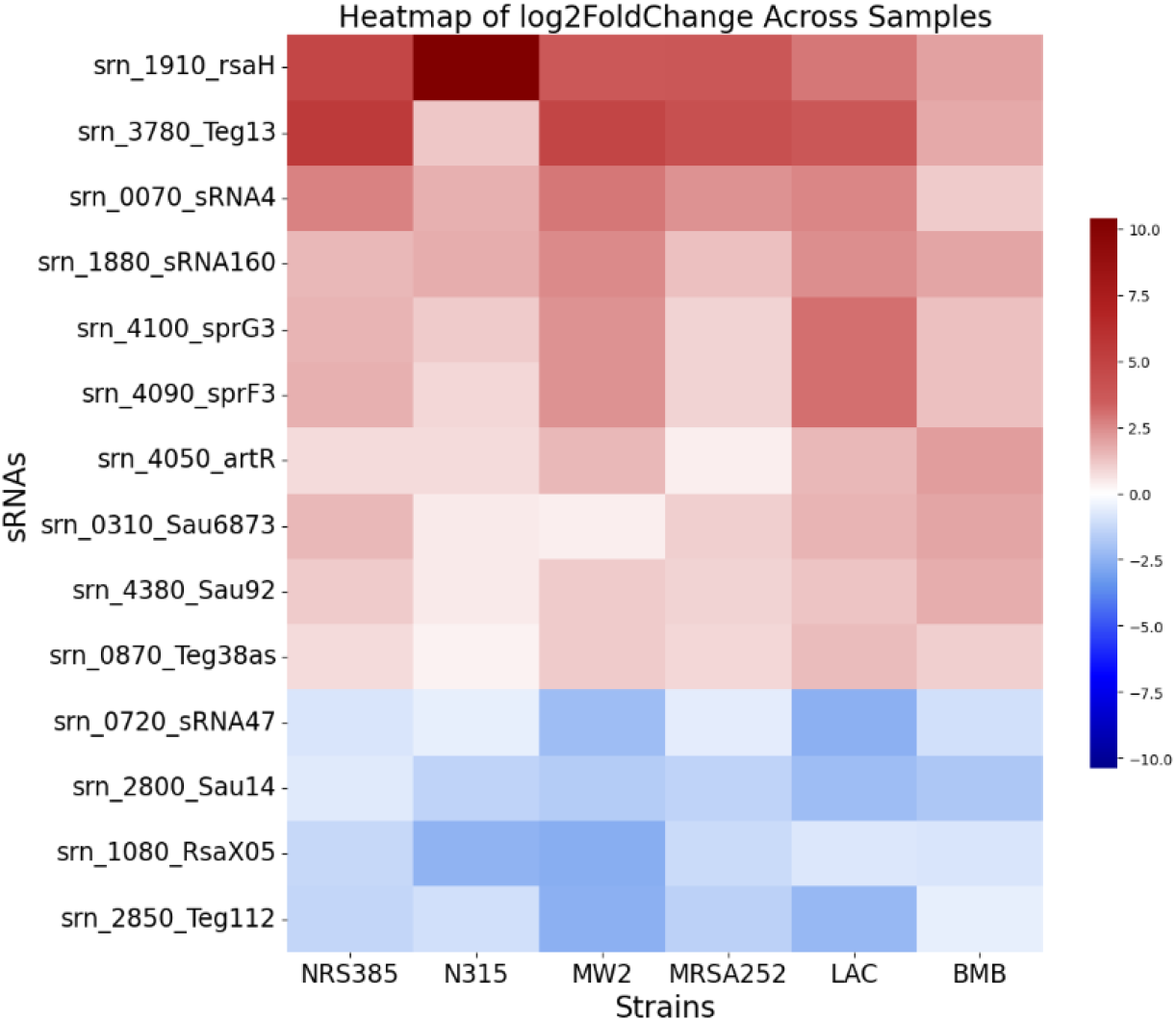
Heatmap of sRNAs consistently up- and down-regulated across all strains. Fourteen sRNAs are highlighted. Those up-regulated include rsaH, Teg13, sRNA4, sRNA160, sprG3, sprF3, artR, Sau6873, Sau92, and Teg38as, while down-regulated sRNAs comprise sRNA47, Sau14, RsaX05, and Teg112. *sRNA’s name according to the most common name entry in the Staphylococcal Regulatory RNAs Database (SRD) (https://srd.genouest.org).

A comprehensive evaluation of upregulated sRNAs under biofilm conditions identified several candidates for deeper investigation. Among 70 sRNAs commonly upregulated across five public *S. aureus* strains, notable examples include rsaH, artR, RsaOV, Teg23, sprF3, sprG3, Teg13, and Sau6667 (Tables S6 to S11). A study by Burke (2018) showed that mutations in these sRNAs altered biofilm phenotypes in a USA300 strain. Notably, rsaH, artR, Teg13, sprG3, and sprF3 were also upregulated in Bmb9393, reinforcing their potential role in biofilm formation and regulation across *S. aureus* lineages (Table S6).

The upregulated sRNAs sprG3 and sprF3 were consistently identified across all analyzed *S. aureus* strains and are particularly notable. These sRNAs are transcribed from genes encoding a toxin-antitoxin (TA) system, in which the antitoxin prevents protein synthesis in trans-acting cells (Chlebicka *et al*., 2021; Sarpong and Murphy, 2021). The SprG3/SprF3 TA systems are implicated in promoting persister cell formation under stress conditions, enhancing biofilm resilience through bacteriostatic activity rather than inducing cell death (Riffaud *et al*., 2019).

The sRNA artR was consistently upregulated and was reported to modulate *hla* expression by repressing *sarT* mRNA, both directly and through RNAIII-mediated degradation. Previous studies showed that *agrA* negatively regulates artR (Schmidt *et al*., 2001; Xue *et al*., 2014); this relationship was reflected in most strains analyzed, except for Bmb9393 and N315, where *agrA* was upregulated under biofilm conditions. The *sarT* expression patterns also varied: it was generally downregulated, whereas artR was upregulated, with slight increases in N315 and NRS385. These results suggest artR may regulate additional targets. Altogether, these findings underscore the complex, strain-specific roles of sRNA-mediated regulation in *S. aureus* biofilm formation.

Prior work proposed that Sau41 represses the Agr circuit by inhibiting RNAIII, providing negative feedback that limits hypervirulence (Yu *et al*., 2023). Here, we specifically hypothesized that the Sau41–RNAIII interaction functions as a competitive sponge, acting as a decoy-like interaction that modulates RNAIII availability, thereby fine-tuning *hla* translation and α-hemolysin output, a configuration compatible with a stable, non-dispersive biofilm architecture (Yu *et al*., 2023). Rather than inhibiting the Agr system, Sau41 could modulate α-hemolysin levels to achieve a balance in which the toxin contributes to biofilm structure by promoting cell-to-cell interactions without dispersing the biofilm. This mechanism aligns with the essential role of Sau41 in biofilm formation, as it dynamically interacts with RNAIII to regulate α-hemolysin expression. In the canonical pathway, RNAIII activates *hla* translation by binding to its 5’UTR, introducing structural rearrangements that expose the SD region to initiate translation (Morfeldt *et al*., 1995). Therefore, as Sau41 levels increase, it competes with *hla* for RNAIII, attenuating RNAIII’s activity (sponge mechanism between the two non-coding RNAs). As previously demonstrated, this results in decreased production of *α*-hemolysin (encoded by *hla*) and *δ*-hemolysin (encoded by RNAIII), which is a secondary function of RNAIII (Yu *et al*., 2023).

Bmb9393, known for producing high levels of biofilm (Costa *et al*., 2023), *hla* (SABB_RS07920; baseMean 154), and RNAIII (baseMean 7,114), displays no significant log2FC between biofilm and planktonic conditions, whereas Sau41 increases (baseMean 3,670; log2FC 1.1) (see Table S6). Although *hla* and RNAIII do not pass the statistical threshold via DEG analysis, the ΔΔCt qRT-PCR profiles recapitulate this pattern. Relative to RNAIII, Sau41 is present at about half the level and *hla* at about twice the level (see ΔΔCt Figure 3). Under a competitive sRNA sponge model, this stoichiometry reduces the free fraction of RNAIII while still permitting activation (and potentially stabilization) of *hla* mRNA, thereby tuning α-hemolysin output. Given Hla’s capacity to lyse immune cells, including macrophages (Scherr *et al*., 2015), such tuning aligns virulence and immune evasion with the maintenance of a cohesive, non-dispersive biofilm, consistent with the Sau41–RNAIII–*hla* axis. The relative stoichiometry of Sau41 and RNAIII may influence the fraction of RNAIII available to activate *hla* translation and thereby affect Hla abundance. By contrast, across the USA-lineage strains examined, RNA-seq profiles for Sau41, RNAIII, and *hla* were heterogeneous. They did not display a coherent stoichiometric arrangement that would parsimoniously explain biofilm behavior under the same model, suggesting additional regulators or lineage-specific network wiring contribute to *hla* control in those backgrounds. Collectively, although Hla is required for biofilm formation (Scherr *et al*., 2015), its expression must be controlled to a specific range, and it likely interacts with other regulatory pathways.

Another important sRNA, rsaA, is exclusively upregulated in the USA strains (Tables S7-S11) and is a well-characterized regulator of biofilm formation (Romilly *et al*., 2014). Romilly et al. (2014) demonstrated that rsaA binds to *mgrA* mRNA, blocking the SD sequence and inhibiting its translation, thereby promoting an alternative, *ica*-independent biofilm pathway. In our data, rsaA was consistently upregulated across all USA strains but not significantly altered in Bmb9393 under biofilm conditions (Tables S6 to S11). The MgrA protein generally mirrors the *agr* program and functions as a negative regulator of biofilm, repressing adhesins and autolysis while activating extracellular enzymes that degrade matrix components (Costa *et al*., 2023). Consequently, *mgrA* downregulation is expected to favor the biofilm state. Consistent with this, Bmb9393 exhibited the most substantial decrease in *mgrA* expression (log2FC −1.42) despite the absence of rsaA upregulation, whereas USA500 (NRS385) showed a moderate decline (log2FC −0.92) alongside rsaA induction (Tables S6 to S11). In contrast, Bmb9393 appears to downshift *mgrA* via alternative regulatory inputs, such as the WalKR two-component system (*walK* was upregulated in Bmb9393; see ΔΔCt in Figure 3), which was suggested to control *mgrA* expression negatively (Guo *et al*., 2025). Both scenarios converge on reduced MgrA output, a configuration that relaxes repression of adhesins and autolysis and limits the activity of matrix-degrading enzymes, thereby reinforcing biofilm maintenance.

### Co-expression modules and sRNA and CDS hubs

The WGCNA co-expression analysis identified 23 modules of co-expressed transcript modules. Seven modules showed significant module-trait correlations with the biofilm condition (*p* ≤ 0.05), while others were associated with the planktonic state (Figure 2). Supplementary Table S12 provides a comprehensive compendium of module assignments for all CDSs and sRNA loci across the studied strains, enabling cross-lineage comparisons and revealing both shared and strain-specific signatures. In WGCNA, modules group transcripts with shared expression dynamics, and hubs are the most centrally connected members, often serving as informative indicators of a module’s underlying biology. In Table S13, we provide details of the hub assignments for each module associated with the planktonic or biofilm state. These modules represent coordinated transcriptional programs in which genes with correlated expression are frequently co-regulated by shared transcription factors or sRNAs. Therefore, these modules can represent distinct physiological states or adaptive responses, providing a functional framework for interpreting the regulatory role of the sRNAs identified as central nodes in the network.

**Figure 2.**
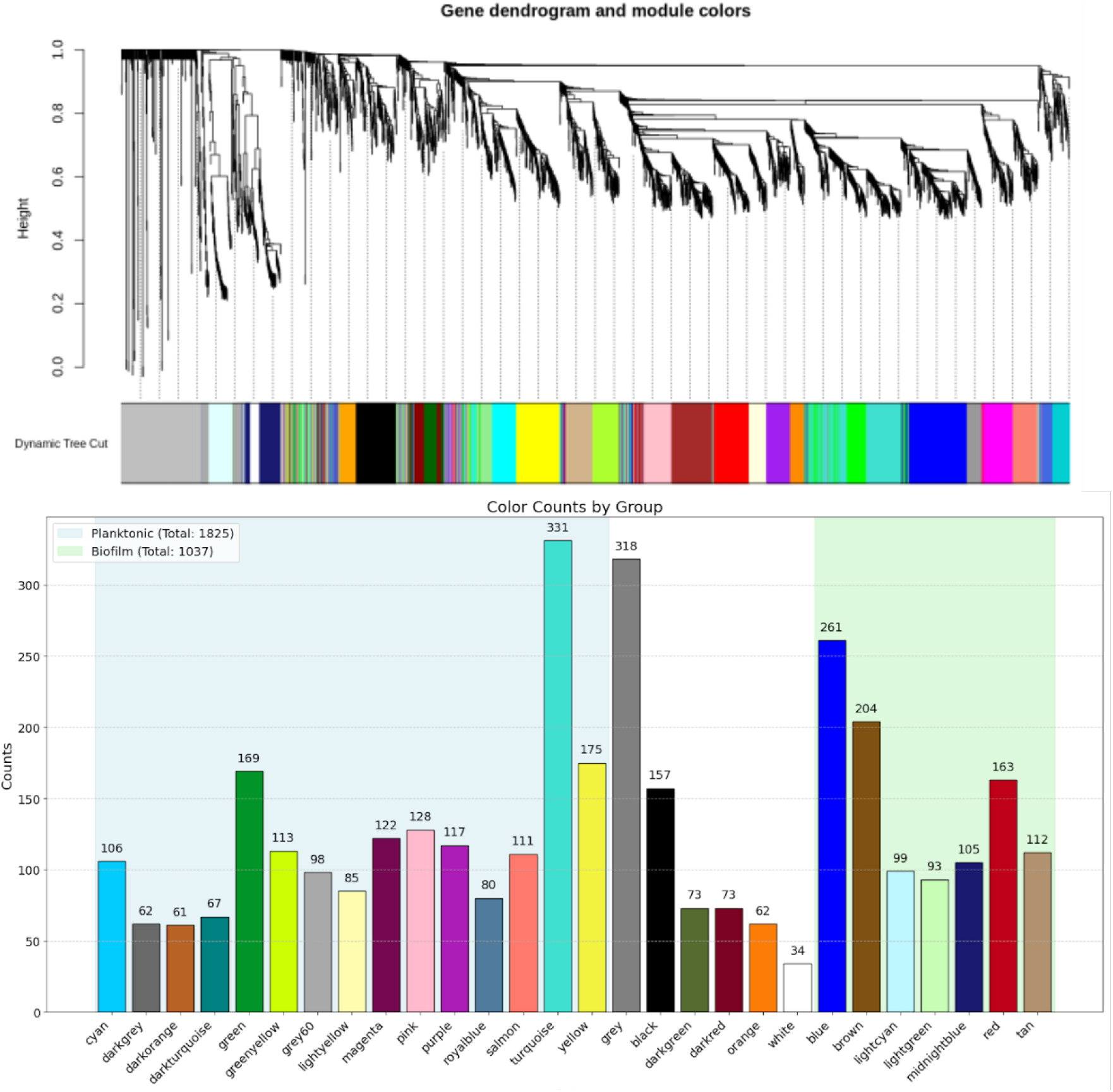
Module Composition in Planktonic and Biofilm Conditions. The top panel shows a hierarchical gene dendrogram produced by WGCNA, where genes are grouped into modules based on co-expression patterns. Each module is assigned a unique color, representing clusters of highly co-expressed genes. The lower panel provides a bar plot summarizing the distribution of module colors across two conditions: planktonic (blue background, total 1,825 genes) and biofilm (green background, total 1,037 genes). Bar heights indicate the number of genes in each module for each condition, revealing differences in module sizes and co-expression patterns between the two environments.

Gene Ontology enrichment yielded biologically interpretable insights for 17 modules (Table S14). Among these, the brown module was the principal biofilm-associated cluster, enriched for metabolic adaptation pathways, including urease activity (*ureA, ureB, ureC*) and nitrogen metabolism (*glnA, narT*), consistent with Tomlinson, Malof, and Shaw (2021). These pathways contribute to pH regulation and survival in nutrient-limited, anaerobic biofilm environments (Park, Choi, and Kim, 2020; Vudhya Gowrisankar *et al*., 2021). The brown module’s hub, SABB_RS04005, encodes a diacylglycerol kinase (DgkB) that recycles diacylglycerol (DAG) for lipoteichoic acid (LTA) biosynthesis, contributing to the cell-envelope homeostasis in *S. aureus* (Miller *et al*., 2008). Perturbations of LTA synthesis, processing, and related signaling pathways have been shown to influence biofilm phenotypes (Ahn *et al*., 2018). Accordingly, the prediction of DgkB as a highly connected node in this biofilm-associated module is consistent with known envelope controls of biofilm phenotype.

The tan module, another biofilm-associated cluster, was enriched with nucleotide metabolism genes (*guaA, guaB*, ppGpp pathways) (Giammarinaro *et al*., 2022), which were downregulated in USA strains but upregulated in Bmb9393 (Tables S6 to S11), suggesting strain-specific adaptations (Giammarinaro et al., 2022). The tan hub gene, SABB_RS03985, encodes an ABC transporter linked to antioxidant defenses via L-ergothioneine (ET) uptake (Y. Zhang *et al*., 2022), with potential roles in ROS/RNS resistance during infection.

Notably, the three components of the Sau41–RNAIII–*hla* axis were localized to modules significantly associated with the biofilm state. Sau41 and RNAIII were co-assigned to the tan module, whereas *hla* mapped to the blue module (Table S12).

Additional hub genes from the modules include SABB_RS14820, an anaerobic ribonucleotide reductase activator involved in low-oxygen adaptation (Ollagnier *et al*., 1997; Ewels *et al*., 2020), and SABB_RS04360, encoding ClfA, a MSCRAMM adhesin critical for host colonization, immune evasion, and endocarditis initiation (Palmqvist *et al*., 2004).

Two particularly notable findings involve sRNAs identified as potential gene hubs in two planktonic-associated modules: srn_1580_sbrC and srn_4120_Teg20. Notably, no targets have yet been validated for these sRNAs. The absence of information linking Teg20 to biofilm or planktonic conditions underscores the novelty of this study, as it represents the first investigation of this association. Teg20, characterized as a *cis*-acting regulator functioning as a riboswitch for *glmS* (Beaume *et al*., 2010; Howden *et al*., 2013). This gene is widely recognized as a ribozyme that catalyzes the conversion of fructose-6-phosphate and glutamine into glutamate and glucosamine-6-phosphate (GlcN6P), which serves as the starting point for bacterial cell wall synthesis (Ferré-D’Amaré, 2010). However, more recent studies involving a *glmS* mutant have elucidated its significant role in *S. aureus* biofilm formation, particularly in the presence of advanced glycation end products (AGEs). These studies describe enhanced biofilm formation facilitated by promoting eDNA release through the upregulation of *sigB* in *S. aureus*. Notably, the *glmS* mutant exhibits several phenotypic changes, including pigment deficiency, reduced hemolytic capacity, downregulation of *hla* and *hld* expression, and the formation of thinner, sparser biofilms.

Furthermore, these findings indicate that, in the mutant strain, *sigB* and biofilm formation are no longer responsive to AGEs (Ni *et al*., 2024). Our DEG analysis showed that *glmS* expression was downregulated under biofilm conditions in most USA strains, which correlates with reduced *hla* expression reported in the previous section (Tables S7 to S11). Notably, *glmS* is part of the green-yellow module, which is associated with planktonic conditions. At the same time, Teg20 showed heterogeneous expression across strains, suggesting that the role of advanced glycation end products in promoting biofilm formation may involve this pathway in a strain-specific manner.

Our findings regarding sbrC showed that this sRNA was significantly downregulated across multiple USA strains in biofilm: MRSA252 (log_2_FC -2.4), NRS385 (log_2_FC -2.9), N315 (log_2_FC -1.2), LAC (log_2_FC -3.3), and MW2 (log_2_FC -3.3) (Tables S7 to S11).Furthermore, sbrC is part of the SigB regulon (Nielsen *et al*., 2011), a critical regulator of stress responses and cellular adaptation. The upregulation of sbrC in planktonic cells suggests its involvement in promoting or maintaining this lifestyle, potentially by regulating factors that inhibit biofilm formation. Its association with the SigB regulon aligns with its proposed role in stress adaptation, a crucial process for survival in the planktonic state. On the other hand, given its regulatory association with SigB, these findings suggest that sbrC may modulate biofilm dynamics in *S. aureus*.

### Computational prediction, integration, and filtering of sRNA-mRNA pairs

For sRNA target prediction, we used the *S. aureus* Bmb9393 reference genome, which we updated with all annotated sRNAs and coding sequences (CDSs). We generated predictions using four established tools —sRNARFTarget, TargetRNA3, IntaRNA, and RNAplex —and applied the thresholds detailed in the materials and methods section. To prioritize interactions supported across programs, we retained pairs that satisfied *p* < 0.1 in TargetRNA3, IntaRNA, and RNAplex and probability ≥ 0.5 in sRNARFTarget. As a direct consequence of these cutoffs, only interactions consistently exceeding these thresholds were carried forward for downstream analyses. To benchmark and justify the cutoffs, we curated 31 literature-supported sRNA–mRNA interactions present in the Bmb9393 locus set as positive controls (Table 3).

Then, we used an in-house–developed Python workflow to aggregate, de-duplicate, and filter the predictions (Figure 6), yielding 1,468 unique sRNA–mRNA pairs involving 139 sRNAs. We further applied network- and expression-consistency filters: we retained pairs only when the sRNA and its putative target were co-assigned to the same WGCNA module and when the direction of DEG (biofilm vs. planktonic) was conserved across the studied strains, permitting at most one discordance of DEG direction. For completeness, the full cross-strain catalog—the combined output of all four prediction programs—is provided in Supplementary Table S15.

A significant gap remains in our understanding of the functions of the majority of the sRNAs under investigation. Specifically, there is limited information on the functions of potential mRNA targets or the pathways they may influence. Furthermore, an even smaller subset has been linked to biofilm formation. We manually selected the most intriguing pairings identified in our analysis, emphasizing their potential significance for biofilm formation. Given the limited knowledge of sRNA function, our discussion primarily centers on the implications of the genes associated with sRNAs. These genes, which play key roles in various cellular and molecular processes, offer valuable insights into biofilm dynamics and regulation. Table 2 provides a comprehensive overview of selected sRNAs discussed in this section, along with their predicted mRNA targets in this study.

**Table 2.**
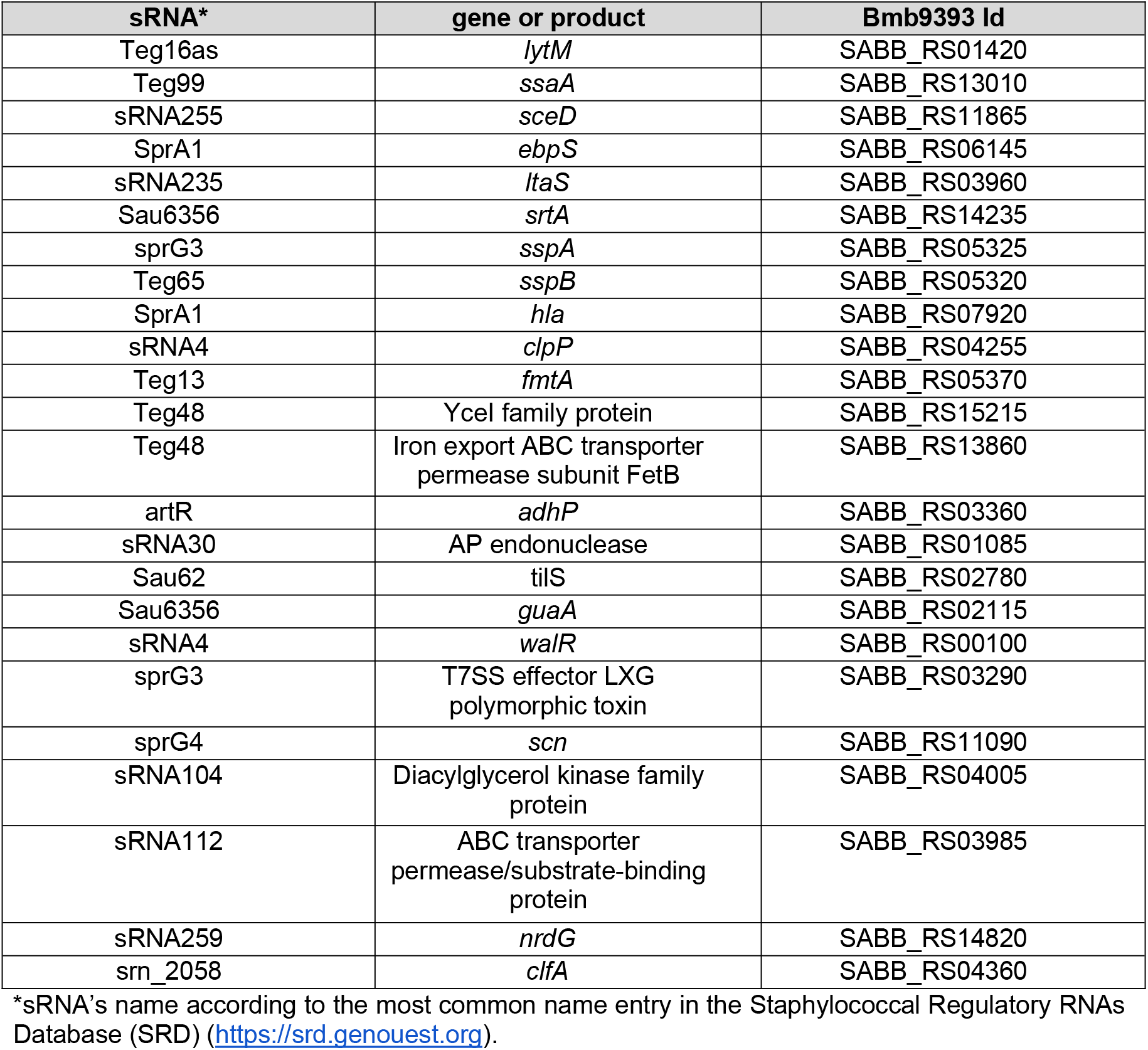
Prioritized sRNAs and their Corresponding Predicted mRNA targets

We first note the SprG4–*scn* interaction: both transcripts are upregulated in biofilm and co-assigned to the brown biofilm module (Tables S12 and S15). The *scn* gene (SABB_RS11090) encodes the *Staphylococcal* complement inhibitor SCIN. This protein is primarily recognized for its role in inhibiting complement activation, a key component of the innate immune response against bacterial infections. SCIN achieves this by binding and stabilizing the human C3 convertases (C4b2a and C3bBb), thereby inactivating them. Once stabilized, the convertases lose their ability to cleave complement C3, thereby preventing further deposition of C3b on the bacterial surface and impeding bacterial phagocytosis. SCIN also inhibits C5a-induced neutrophil responses (Rooijakkers *et al*., 2005). The biofilm itself provides an additional critical immune-evasion mechanism. Together, SCIN and the biofilm could synergistically hinder the immune system’s ability to mount an effective response (Resch *et al*., 2005). This pair represents a promising candidate for future experimental studies.

The predicted pairing between sRNA sprG3 and the mRNA encoding a T7SS LXG polymorphic toxin (SABB_RS03290), both downregulated in biofilm and co-assigned to the lightyellow module (Tables S12 and S15), points to a possible connection with virulence pathways, consistent with the implication of T7SS in *S. aureus* virulence. The encoded protein carries an N-terminal LXG domain, a hallmark of T7SS substrates. Several characterized LXG proteins act as toxins and are co-produced with cognate antitoxin factors, typically encoded immediately downstream. Such antagonistic systems can foster kin discrimination and spatial structuring within biofilms, potentially limiting conflict among cohabiting strains (Bowman and Palmer, 2021). Determining whether sprG3 modulates this locus, and how its expression balance shifts across conditions, may clarify how interbacterial antagonism is coupled to persistence in the biofilm niche.

The pair formed by the sRNA artR and *adhP* mRNA (SABB_RS03360) was upregulated under biofilm conditions (Table S15). Interestingly, *adhP* was also significantly upregulated in the biofilm of *Stenotrophomonas maltophilia*, as reported by Alio et al. (2020). In *E. coli, adhP* has been described as an ethanol-inducible dehydrogenase Thomas *et al*., 2013. Alio et al. (2020) proposed that *adhP* likely contributes to the production of propanol or other short-chain alcohols under biofilm conditions (Alio *et al*., 2020). However, its role in biofilm development or maintenance remains unexplored. Further analysis of *adhP* under the biofilm conditions of *S. aureus* Bmb9393 could provide valuable insights into its functional contributions to biofilm formation and dynamics.

We detected high-confidence interactions between sRNAs and hub coding genes within four biofilm-associated WGCNA modules (Table S13), revealing module-specific coupling of adhesion, metabolism, lipid signaling, and transport to sRNA control. In the blue module, *clfA* (SABB_RS04360) is predicted to be targeted by srn_2058; in the red module, sRNA259 aligns with *nrdG* mRNA (SABB_RS14820; class III anaerobic ribonucleotide reductase); in the brown module, sRNA104 pairs with a diacylglycerol kinase–family protein (SABB_RS04005); and in the tan module, sRNA11 associates with the mRNA for an ABC transporter permease/substrate-binding protein (SABB_RS03985). Together, these associations, retained after cross-tool, co-expression, and expression-consistency filters, underscore coordinated sRNA regulation across biofilm-relevant functional hubs. Future experimental validation of these interactions will be essential to uncover the precise mechanisms by which sRNAs influence biofilm dynamics in *S. aureus*.

### Genomic context for constraining sRNA target prediction

The genetic location of sRNAs is critical in target prediction, especially for antisense sRNAs transcribed in the opposite direction of a CDS. In such cases, predicting the corresponding CDS as the target gene is straightforward due to their complementary orientation. This natural complementarity allows for perfect or near-perfect base pairing, which significantly enhances the likelihood of interaction, for example, between an antisense sRNA and its target gene (Georg and Hess, 2011). Then, we annotated the Bmb9393 genomic context of each sRNA (e.g., intergenic, intragenic, UTR-associated, antisense) and the associated CDS in Table S15, indicating whether the sRNA is in the sense or antisense orientation.

In this context, four sRNAs predicted as antisense in Bmb9393 can be highlighted: sRNA30, Sau6356, Sau62, and sRNA4. We finally filtered their corresponding targets with high confidence on the opposite and complementary strands, specifically SABB_RS01085 (sRNA30), SABB_RS02115 (Sau6356), SABB_RS02780 (Sau62), and SABB_RS00100 (sRNA4).

Among these predicted antisense sRNAs, a particularly interesting finding is the interaction between sRNA4 and *walR* (SABB_RS00100), both of which are co-expressed in the turquoise module associated with the planktonic state (Tables S12 and S13). Notably, sRNA4 was one of the 10 sRNAs consistently upregulated across all six strains analyzed in this study, including the Bmb9393 strain (Table S6 and confirmed by qRT-PCR, Figure 3). This consistent expression underscores its potential regulatory importance and suggests a conserved role in biofilm-associated processes.

**Figure 3.**
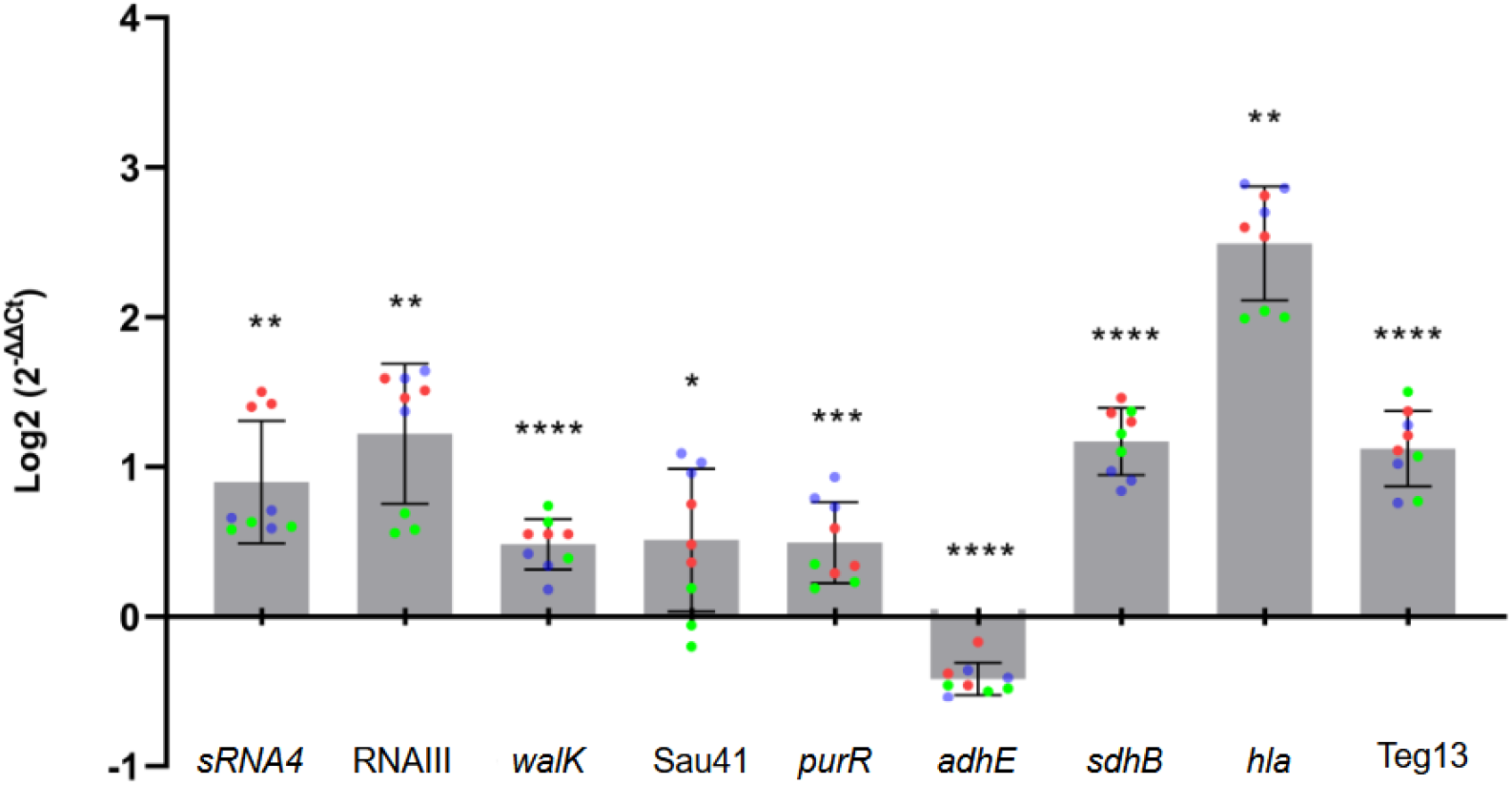
Real-Time qRT-PCR Validation of sRNA and mRNA Expression Under Biofilm Conditions (Bmb9393). qRT-PCR (ΔΔCt) comparison of biofilm versus planktonic conditions for nine loci in Bmb9393: *walK* (SAB_RS00105), sRNA4, RNAIII, sau41, *hla, purR* (SABB_RS02705), *adhE* (SABB_RS00775), *sdhB* (SABB_RS08000), and Teg13. Relative expression was computed by ΔΔCt (see Methods for calculation and sign convention). The direction of change (up/down in biofilm) matches the RNA-seq results for all loci (see Supplementary Table S6). sRNA’s name according to the most common name entry in the Staphylococcal Regulatory RNAs Database (SRD) (https://srd.genouest.org). The colored dots on the bars denote the three technical replicates (same color) for each of the three biological replicates (different colors).

WalR, also upregulated in biofilm (Table S15), is the response regulator of the WalRK two-component system, a central controller of cell-wall metabolism in *S. aureus* (Fabret and Hoch, 1998; Dubrac *et al*., 2007; Costa *et al*., 2023). This system is essential in *Firmicutes* and is highly conserved among low GC% Gram-positive bacteria, including pathogens such as *Streptococcus pneumoniae* and *S. aureus* (Takada and Yoshikawa, 2018). Validated targets of the WalR regulon include genes involved in cell wall degradation and autolysis, such as *lytM, ssaA*, and *sceD*, as well as the gene encoding the adhesin EbpS linked to virulence (Dubrac *et al*., 2007; Delauné *et al*., 2012). One of its most significant targets is *ltaS*, which encodes lipoteichoic acid (LTA). As mentioned before, LTA is crucial to the cell-envelope homeostasis in *S. aureus* (Gründling and Schneewind, 2007; Miller *et al*., 2008). The essentiality of *ltaS* has been demonstrated in multiple independent studies, with *ltaS* mutants found to be inviable (Chaudhuri *et al*., 2009; Santiago *et al*., 2015; Chen *et al*., 2017). This observed consistency suggests a direct connection between *ltaS* and the essential role of the WalRK system in biofilm (Delauné *et al*., 2012; Costa *et al*., 2023). Additionally, overexpression of *ltaS* enhances autolysis, hemolytic activity, and the transcription of virulence genes associated with host interaction, cytolysis, and immune evasion, partly through activation of the TCS SaeSR (Delauné *et al*., 2012).

Beyond its role in biofilm regulation, WalRK has also been associated with antimicrobial resistance, particularly vancomycin resistance. Strains with intermediate resistance to vancomycin frequently harbor mutations in the *walRK* promoter and operon genes (Hafer *et al*., 2012; Peng *et al*., 2016). Notably, *ltaS* is an appealing drug target because LTA is absent in eukaryotic cells (Gründling and Schneewind, 2007).

In contrast to *Enterococcus faecalis*, where an antisense sRNA targeting *walR* reduces WalR and biofilm formation (Wu et al., 2020), the concordant increase in sRNA4 and *walR* in *S. aureus* suggests a stabilizing role for *walR*. We observed that sRNA4 (minus strand, nt 24,953–25,188) lies antisense to *walR* (plus strand, nt 24,903–25,604; SABB_RS00100), creating a CDS-internal overlap. Both transcripts are upregulated in biofilm, and qRT-PCR corroborates the upregulation of sRNA4. We hypothesize that this internal antisense pairing either occludes endonucleolytic sites and limits ribonuclease access or functions as a post-transcriptional fine-tuner under biofilm conditions to sustain *walR* when transcriptional input is high, both mechanisms consistent with elevated steady-state *walR* despite duplex formation. This is a compelling hypothesis for future experimental testing.

On the other hand, intragenic (sense) sRNAs are unlikely to base-pair with the mRNAs of their host genes because sense–sense pairing lacks the complementarity required for canonical sRNA–mRNA regulation. Accordingly, these sRNAs are often co-transcribed with their hosts and tend to mirror their expression profiles. Likewise, sRNAs positioned in 5′ or 3′ UTRs (distinct from bona fide riboswitches) do not inherently have a higher probability of targeting their cognate genes (Wagner and Romby, 2015).

Among the intragenic sRNAs, Teg13 (encoded within SABB_RS11020) is one of ten consistently upregulated across all six strains (Tables S6 to S11), and as mentioned before, its gene deletion causes a defective biofilm phenotype. After applying our multi-stage filters, we retained 58 candidate Teg13 target pairs, all of which grouped to the yellow (planktonic-associated) module (Table S15). Notably, Teg13 and its host gene SABB_RS11020 were co-assigned to the yellow module (i.e., co-expressed) (Table S12). However, our pipeline did not predict a Teg13–SABB_RS11020 interaction, consistent with expectations for intragenic (sense) sRNAs.

Among Teg13’s predicted targets, fmtA, a well-characterized modulator of methicillin resistance in *S. aureus*, is especially notable. Analysis of the *fmtA* promoter demonstrated that the transcriptional level of *fmtA* increased when the cells were exposed to β-lactam antibiotics (Komatsuzawa *et al*., 1999). In our data, *fmtA* is downregulated in biofilm across the USA-lineage strains (but unchanged in Bmb9393). In comparison, Teg13 is upregulated in biofilm and co-assigned with its host gene to the yellow module. This pattern supports the hypothesis that Teg13 contributes to post-transcriptional repression of *fmtA* in a lineage-dependent manner. Also, previous studies have demonstrated that *fmtA* expression is regulated by SarA, a critical regulon involved in biofilm formation (Komatsuzawa *et al*., 1999). SarA promotes biofilm development by negatively regulating protease and nuclease activities, essential for biofilm maintenance and stability (Abdelhady *et al*., 2014; Atwood *et al*., 2015; Liu *et al*., 2024). Biologically, decreasing FmtA during biofilm growth could sensitize cells to β-lactams, a potentially favorable therapeutic outcome. However, it may also increase autolysis and eDNA release, effects that reinforce matrix integrity and stabilize the biofilm, thereby complicating eradication. These observations argue for context-specific strategies: if Teg13-mediated repression of *fmtA* is confirmed, combining Teg13 overexpression with β-lactams and matrix-disrupting adjuncts (e.g., DNase or anti-biofilm agents) could enhance β-lactam susceptibility while limiting biofilm matrix reinforcement.

As an example of target prediction for an sRNA associated with the UTR region of a CDS, we highlight the neglected Teg48, derived from the 5’ UTR of SABB_RS03420 (Tables S1 and S15). Three possible mRNA targets were identified: SABB_RS13860 (iron export ABC transporter permease subunit FetB), SABB_RS15215 (YceI family protein), and SABB_RS16565 (Uncharacterized protein) (Table S15).

Regarding Teg48 and *fetB*, our DEG analysis revealed consistent upregulation of both loci across all evaluated strains (Tables S6-S11). Interestingly, Nicolaou *et al*. (2013) characterized the ABC iron transporter system FetAB and demonstrated its role in enhancing resistance to H2O2-mediated oxidative stress. They also showed that FetAB mitigates oxidative stress under iron overload conditions by facilitating iron homeostasis. Our observed upregulation of *fetB* and Teg48 in *S. aureus* biofilm conditions across all strains suggests a potential protective mechanism against oxidative stress, promoting iron export and maintaining iron homeostasis. To further elucidate the functional role of this regulatory relationship in *S. aureus* biofilm biology, experimental validation is essential. Such studies could include assessing ROS levels, evaluating biofilm viability under oxidative stress, and performing knockdown analyses of Teg48 and *fetB* to determine their independent and combined contributions.

### qRT-PCR validation of sRNA and coding-gene expression under biofilm conditions

To orthogonally validate our RNA-seq calls, we quantified transcript levels by qRT-PCR (ΔΔCt) for nine loci for strain BMB9393: *walK* (SAB_RS00105), sRNA4, RNAIII, Sau41, *hla, purR* (SABB_RS02705), *adhE* (SABB_RS00775), *sdhB* (SABB_RS08000), and Teg13, comparing biofilm with planktonic conditions. The ΔΔCt profiles recapitulated the direction of change (upregulation vs downregulation) observed by RNA-seq across all loci, with expected differences in magnitude owing to platform-specific dynamic ranges and normalization (see ΔΔCt Figure 3 for qRT-PCR and supplementary log2FC Table S6 for RNA-seq).

Biologically, the qRT-PCR direction reinforces the regulatory model from our network analyses. The Sau41–RNAIII–*hla* axis behaves consistently with a sponge-mediated titration model, in which Sau41 availability modulates RNAIII’s capacity to activate *hla* translation and tunes α-hemolysin output in biofilm.

Changes in *walK* align with envelope-remodeling demands typical of sessile growth, while *purR* shifts are compatible with the nucleotide-metabolism reprogramming. Adjustments in *sdhB* and *adhE* hold together with metabolic adaptation to biofilm microenvironments (enhanced fermentative flux and altered TCA activity).

Finally, sRNA4 and Teg13 exhibited concordant qRT-PCR and RNA-seq directions and, as noted earlier, were upregulated in biofilm across all six strains, supporting their prioritization and inferred roles within the predicted regulatory network (see ΔΔCt Figure 3 and supplementary log2FC Table S6).

## Conclusions

We present the first genome-wide annotation of non-coding sRNAs in the *S. aureus* ST239 Bmb9393 strain and a standardized cross-lineage catalog (ST239 and USA lineages), providing a starting point for comparative sRNA analyses and target discovery. Differential expression between biofilm and planktonic states revealed both conserved and lineage-specific programs: 10 sRNAs were consistently upregulated, and four were consistently downregulated across all six strains. Co-expression analysis identified 23 modules, seven of which were significantly associated with biofilm. Within these biofilm-linked modules, Sau41 and RNAIII were co-assigned, a pattern compatible with a competition-based molecular-sponge model between the two sRNAs in biofilm. We identified high-priority interactions between sRNAs and mRNAs at functional hubs, including sRNA2058–*clfA* (adhesion), sRNA259–*nrdG* (anaerobic DNA synthesis), sRNA104–*dgkB* (lipid/LTA signaling), and sRNA11–ABC transporter (stress/transport), each of which is assigned to biofilm-associated modules. Our in-house integrative pipeline (four predictors + module co-assignment + cross-strain DEG-direction consistency) reduced >1.7 million raw calls to 1,468 high-confidence sRNA–mRNA pairs involving 139 sRNAs; we release this curated, machine-readable dataset to guide hypothesis-driven validation.

Collectively, this study delivers a ranked, cross-lineage map of sRNA circuitry in *S. aureus*, a reproducible, evidence-weighted filtering strategy, and experimentally tractable hypotheses linking sRNAs to biofilm stability, envelope homeostasis, and antimicrobial response. The accompanying resources are reusable and FAIR-aligned, enabling extension to additional strains, conditions, and bacterial species.

## Methods

To characterize small RNAs (sRNAs) and their targets in *S. aureus*, with an emphasis on prioritizing high-confidence sRNA–mRNA interactions, we followed four steps: (i) identified and curated sRNAs in the Bmb9393 strain and in additional strains with incomplete ncRNA annotations; (ii) analyzed RNA-seq data from planktonic and biofilm conditions using differential expression (DE) and weighted gene co-expression network analysis (WGCNA); (iii) predicted sRNA–mRNA interactions on the Bmb9393 reference genome; and (iv) applied an in-house, customized multi-criterion filter to retain high-confidence pairs. Figure 4 summarizes the workflow; full details are provided below.

**Figure 4.**
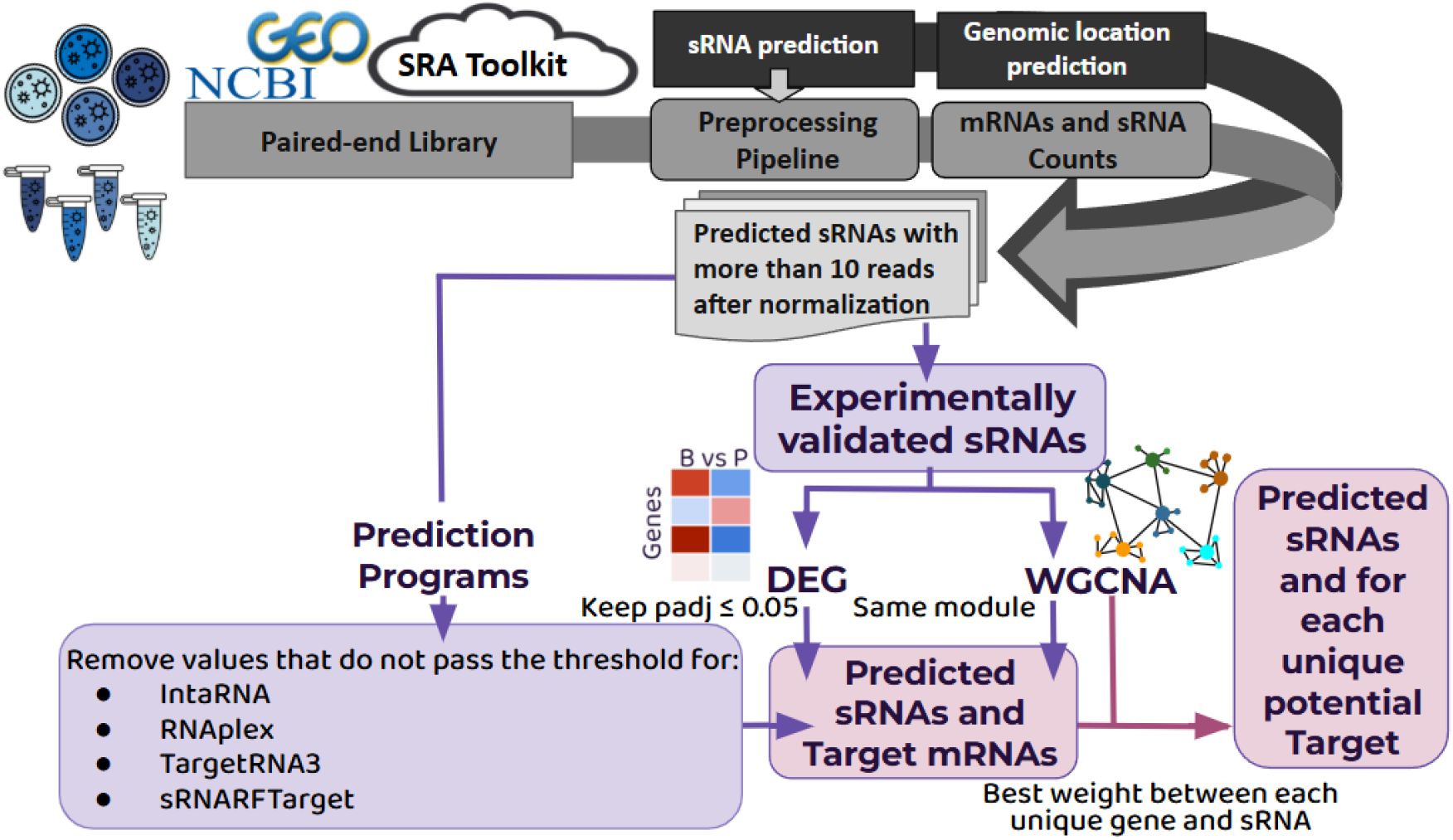
Workflow for characterizing small RNAs (sRNAs) and prioritizing sRNA–mRNA interactions in *S. aureus*. (1) Data acquisition: paired-end RNA-seq (in-house and GEO/SRA) and genomes for Bmb9393, N315, MRSA252, LAC/USA300 FPR3757, and MW2 (Table S16). (2) sRNA prediction & annotation: strand-aware prediction and classification by genomic context (intergenic, antisense, UTR). (3) RNA-seq processing (Bmb9393): in-house pipeline from raw reads to gene-level counts, normalization, and differential expression (biofilm vs planktonic). (4) Target prediction: *in silico* identification of candidate sRNA targets. (5) Network integration: co-expression (WGCNA) clusters genes/sRNAs; pairs co-assigned to the same module are flagged as putative regulatory links. (6) Prioritization: applying thresholds and network/expression filters reduces the initial set to high-priority candidates for downstream experimental validation.

### Identification of sRNAs

To identify sRNA in this study, two distinct approaches were employed. The first method relied on sequence homology detection using the non-redundant sequences available at the NR2 (https://nr2.ncrnadatabases.org) and SRD (Sassi *et al*., 2015) databases. The second approach involved predicting and annotating sRNAs using the StructRNAFinder pipeline, based on secondary-structure covariance models derived from known RNA families (Arias-Carrasco *et al*., 2018). Results from the two approaches (sequence homology and covariance model predictions) were combined for consolidation. For consistency, sRNAs were named according to the most common name entry in the Staphylococcal Regulatory RNAs Database (SRD) (https://srd.genouest.org).

Subsequently, we classified each predicted sRNA by genomic context relative to annotated genes in the study genomes: Bmb9393, N315, MRSA252, LAC (USA300/FPR3757), and MW2. Because a well-annotated reference for NRS385 (USA500) is unavailable, we used the LAC (USA300/FPR3757) genome as a proxy for USA500, consistent with prior studies (Li *et al*., 2009; Glaser *et al*., 2016; Tomlinson, Malof, and Shaw, 2021). The sRNAs were categorized into five genomic locations: (a) antisense: located on the opposite strand of a CDS, (b) intragenic: situated within a CDS on the same strand, (c) 5’ UTR: located upstream of a gene, (d) 3’ UTR: located downstream of a gene, and (e) intergenic: situated between genes outside the defined UTR regions.

### Bacterial growth under biofilm and planktonic conditions

For biofilm formation, the Bmb9393 strain was streaked on a TSA plate and incubated for 24 h at 37 °C. Approximately three colonies were inoculated into 2 mL of TSB supplemented with 1% (w/v) glucose (TSB-Glu) and incubated for 24 h at 37 °C with shaking at 250 rpm. The resulting cultures were diluted 1:100 in TSB-Glu, and 200 μL aliquots were dispensed into 48 wells of sterile 96-well plates (Nunclon untreated; Nunc A/S, Roskilde, Denmark), followed by a 24-h incubation at 37 °C. After removal of supernatants, biofilms were washed with sterile water to remove non-adherent cells. Biofilms were then recovered by scraping with sterile toothpicks into 200 μL of TSB-Glu and immediately treated with an equal volume of cold acetone–ethanol solution (1:1, v/v). Samples were stored at −80 °C until RNA preparation. For planktonic cell preparation, 18-h cultures of BMB9393 were diluted 1:100 in TSB-Glu and incubated for 24 h at 37 °C with shaking at 250 rpm. Cells were harvested, mixed with an equal volume of cold acetone– ethanol solution (1:1, v/v), and stored at −80 °C until RNA preparation. A qRT-PCR experiment was performed in biological triplicates, with three technical replicates per condition. While the RNA-seq experiment was performed in biological independent quadruplicates (Ferreira *et al*., 2012; Beltrame *et al*., 2015; Lade *et al*., 2019).

### RNA extraction

Frozen bacterial cell suspensions in acetone–ethanol solution were thawed on ice and centrifuged at 8,700 × g at 4 °C. The pellet was washed twice with TES buffer (20 mM Tris-HCl, pH 7.6, containing 10 mM EDTA, pH 8.0, 50 mM NaCl, and 20% wt/vol sucrose), using half the original volume of the cell suspension, followed by centrifugation. Bacterial cells were adjusted to an optical density of 0.8 at 600 nm (approximately 108–10^9^ CFU/mL) using a SpectroMax Plus 384 spectrophotometer (Molecular Devices, Silicon Valley, CA, USA). The bacterial pellet was then resuspended in 10 μL of a 10 U/μL lysostaphin solution and 90 μL TES buffer. The mixture was incubated on ice for 30 min, followed by 5 min at 37 °C (Beltrame et al., 2015). After bacterial lysis, RNA extraction was performed using the RNeasy Mini Kit protocol (Qiagen, Hilden, DE, Germany). The quantity of extracted RNA was measured with a Nanodrop spectrophotometer (Thermo Fisher Scientific, Waltham, MA, USA). A standard PCR amplification was performed with *rrs* primers (Table 1) to ensure the absence of DNA in the RNA preparations. The integrity of total RNA was initially assessed by visualizing the 23S/16S rRNA banding pattern using 1.2% agarose gel electrophoresis in 1× TAE buffer (20 mM Tris-acetate, 0.5 mM EDTA, pH 8.0) run at 110 V for 50 min. The gel was stained with ethidium bromide and visualized using a gel documentation system (Gel-Doc EZ Imager, BioRad, Hercules, CA, USA).

### RNA-Seq analysis of paired-end libraries

The Qiagen Stranded RNA Library Kit, which preserves strand-specific information and enables simultaneous capture of both sRNA and mRNA transcripts, was used to construct the libraries of the *S. aureus* Bmb9393 strain. Next, sequencing was performed on the Illumina NextSeq 500/550 using the NextSeq 500/550 v2 kit (75 cycles) at the UGCDFA of the Laboratório Nacional de Computação Científica (https://www.labinfo.lncc.br/ugc), Brazil.

Additionally, we analyzed the paired-end data from a large-scale RNA expression profiling experimentation (Tomlinson, Malof and Shaw, 2021), which conducted a comprehensive transcriptomic analysis of five MRSA strains: N315 (USA100), MRSA252 (USA200), LAC or USA300 FPR3757 (USA300), MW2 (USA400), and NRS385 (USA500) (Tomlinson, Malof and Shaw, 2021) (see dataset description in Table S16). To perform a comparative analysis with the Bmb9393 strain, we retrieved data from the NCBI Gene Expression Omnibus (GEO) (Edgar, 2002) for the 24-hour growth time point in both conditions using the SRA Toolkit (https://github.com/ncbi/sra-tools). The quality of the raw reads was first evaluated with FastQC, followed by adapter trimming and quality filtering with FastP (Chen *et al*., 2018). The quality of the processed reads was then reassessed using FastQC to ensure data integrity (https://github.com/s-andrews/FastQC). High-quality reads were aligned to the reference genome provided in FASTA format and annotated with a GTF file using Rsubread (Liao, Smyth, and Shi, 2019). The alignment files were subsequently processed with SAMtools (Danecek *et al*., 2021), including conversion to BAM format, sorting, and indexing. Finally, gene-level expression was quantified using FeatureCounts (Liao, Smyth, and Shi, 2014), which incorporated genome annotations to generate a read count matrix for downstream analyses. The raw count matrix generated during preprocessing was used for Differential Expression Analysis between biofilm and planktonic conditions using the nf-core/differential-abundance pipeline (Ewels *et al*., 2020).

### Real-Time qRT-PCR

Transcripts were quantified using the real-time qRT-PCR method and the Power SYBR Green RNA-to-CT™ 1-Step kit (Applied Biosystems, Foster City, CA, USA), following the manufacturer’s instructions, on a StepOne Real-Time PCR System (Applied Biosystems, São Paulo, SP, Brazil). The *rrs* transcript encoding 16S rRNA served as an endogenous control. Results were normalized, with the calibration sample assigned a value of 1.0. Primers were designed using Beacon Designer for SYBR® Green (https://www.premierbiosoft.com/molecular_beacons/) and are listed in Table S17. Gene expression data (2−ΔΔCt) were log_2_-transformed before statistical analysis, with the reference group (1) set to 0. Normality was assessed using the Shapiro–Wilk test, followed by either a one-sample t-test or a Wilcoxon signed-rank test against 0, as appropriate. Results are presented as mean ± SD of log_2_-transformed values.

### Construction and analysis of the co-expression network using WGCNA

The count matrix generated during preprocessing was normalized to ensure accurate and reliable downstream analyses. The TMM (Trimmed Mean of M values) normalization method (Robinson and Oshlack, 2010) was applied to correct compositional differences across samples in the first step. After adjusting library sizes and reducing technical biases with TMM, RPKM (Reads Per Kilobase per Million mapped reads) normalization was performed. This additional step accounted for gene length and sequencing depth, providing a more balanced expression profile (Zhao, Ye, and Stanton, 2020). The entire normalization approach was executed in R using the edgeR (4.4.1) package (Chen *et al*., 2024). To ensure the robustness of the results and minimize batch effects from different experiments, an additional analysis was executed using the *limma* package (v3.62.1) (https://bioconductor.org/packages/limma) in R. Genes with zero counts in more than 75% of the samples were excluded to avoid biases from low-expression genes.

The resulting expression matrix was used as input for WGCNA (Langfelder and Horvath, 2008). To mitigate potential biases introduced by excessively high expression values, a log2 transformation was applied, resulting in a more balanced distribution (Zhu *et al*., 2022). The co-expression network was constructed by calculating an adjacency matrix that quantifies the connection strength between genes based on expression similarity. A crucial parameter in this process was the selection of a soft threshold power of 4, which ensured the network’s scale-free topology, a key property for maintaining a power-law distribution of gene connection degrees. The adjacency matrix was then converted into a TOM, which captures direct and indirect connections among genes and provides a robust view of co-expression networks (Langfelder and Horvath, 2008; Soberanes-Gutiérrez *et al*., 2022).

Hierarchical clustering was performed on the TOM to group genes into modules based on their topological overlap. Distinct modules were identified using the dynamic tree-cut method applied to the resulting dendrogram. This method precisely identified co-expressed gene modules, representing groups of genes with highly correlated expression patterns. Module eigengenes were calculated to summarize the expression profiles of genes within each module. These eigengenes facilitated correlations with external traits or phenotypes of interest, such as biofilm and planktonic conditions (Langfelder and Horvath, 2008; Soberanes-Gutiérrez *et al*., 2022).

The biological functions of significant modules were inferred using Gene Ontology (GO) enrichment analysis of the genes within each module. This analysis identified associated biological processes, molecular functions, and cellular components, offering detailed insights into the roles of genes within each module (Soberanes-Gutiérrez *et al*., 2022). For GO enrichment, the N315 strain was selected due to its superior annotation in UniProt, with 2,583 entries in UniProtKB (https://www.uniprot.org/taxonomy/158879). Using the ClueGO app (Bindea *et al*., 2009) in Cytoscape (Shannon *et al*., 2003), modules were enriched individually for GO terms. Only terms with a p-value below 0.05 were retained. The resulting spreadsheets for each module were consolidated into a single document and used for visualization.

To identify the top hub genes within each module, the chooseTopHubInEachModule function from the WGCNA package was employed (Langfelder and Horvath, 2008). This function determines the most connected gene in each module based on intramodular connectivity, which reflects a gene’s correlation with other genes in the same module. Modules labeled as “grey,” representing unassigned or noise-associated genes, were excluded from the analysis to focus on biologically relevant groups. After identifying the top hub genes, each was individually searched in the UniProt database to retrieve detailed functional annotations. Additionally, GO terms were collected using QuickGO (Binns *et al*., 2009).

### Computational prediction, integration, and filtering of sRNA-mRNA pairs

To generate initial sRNA–mRNA interaction calls, we utilized four well-established prediction programs: IntaRNA (v2.0) (Mann, Wright, and Backofen, 2017), TargetRNA3 (v3.0) (Tjaden, 2023), RNAplex (v2.7) (Tafer and Hofacker, 2008), and sRNARFTarget (v1.0) (Naskulwar and Peña-Castillo, 2022). Each program received an input file listing sRNAs with more than 10 reads after TMM normalization, followed by RPKM normalization in Bmb9393. Only sRNAs meeting this threshold were considered successfully predicted in the Bmb9393 strain, serving as the first elimination criterion.

All programs were executed with their default parameters, except for IntaRNA, which offers multiple “personalities” —pre-configured models tailored for specific approaches. In this study, we employed the IntaRNAsTar module, optimized for large-scale, genome-wide target predictions (Tieng *et al*., 2023).

The prediction programs produced a substantial number of results, representing all possible combinations of sRNAs and targets evaluated by the prediction tools, without any filtering. Due to the volume of data, an in-house Python script was developed to refine the results into a more focused selection of potential sRNA-target pairs.

To calibrate tool-specific thresholds, we curated 31 literature-supported sRNA– mRNA interactions present in the Bmb9393 locus set as positive controls. We selected program-specific score cutoffs that maximized recovery of these positives while limiting spurious calls. These cutoff thresholds were established based on the lowest p-value predicted from IntaRNA (0.41), RNAplex (0.76), and TargetRNAIII (0.89), and the weakest probabilities predicted by sRNARFTarget (0.45) (Table S18). Pairs that did not meet the tool-specific criteria were discarded.

Subsequently, DEG analysis was used as a filter to identify sRNAs and their corresponding targets exhibiting significant changes in expression. The main criterion for selection was identifying sRNAs with significant expression (*padj* ≥ 0.05) in at least five of six strains (allowing at most one discordant strain with DEG).

Additionally, using WGCNA, we retained pairs in which the sRNA and its putative mRNA target co-assigned to the same module, and, when multiple sRNAs targeted the same gene, we prioritized the sRNA with the highest intramodular connectivity/weight. Finally, we annotated genomic context (intergenic, intragenic, UTR-associated, antisense) in Bmb9393 to aid downstream prioritization for experimental validation.

## Code and data availability

The scripts used in this study are publicly available at the project’s GitHub repository: https://github.com/carolina5massena/Integrative-RNA-Seq-and-Co-expression-Analyses-Uncover-Confident-sRNA.git. Raw RNA-seq data have been deposited in the NCBI SRA with the Accession SUB15633288, under the BioProject PRJNA1327938, and made public upon publication. This link provides access for editors and reviewers:
https://dataview.ncbi.nlm.nih.gov/object/PRJNA1327938?reviewer=v0k7r57s61raomc4229vc2lto2 to use during the peer review.

## Acknowledgments

This work was supported in part by Conselho Nacional de Desenvolvimento Científico e Tecnológico (CNPq) grant # 305895/2022-2 to MFN and grants # 408725/2022-2 and 306942/2023-2 to AMSF; Fundação Carlos Chagas Filho de Amparo à Pesquisa do Estado do Rio de Janeiro (FAPERJ) grant # E-26/200.555/2023 to MFN and grants # E-26/211.554/2019; E-26/210.064/2020 and E-26/203.941/2024 to AMSF; Coordenação de Aperfeiçoamento de Pessoal de Nível Superior (CAPES) grant # 88881.004652/2024-01 to MFN, and Agencia Nacional de Investigación y Desarrollo de Chile (ANID) grant # 11251012 to JEMH. The authors thank Alexandra Lehmkuhl Gerber and Ana Paula Campos Guimarães for generating the sequences of strain Bmb9393 at UGC/LNCC.

## Ethics Statement

This study involved no animal or human experiments. There are no ethical issues involved.

## Author Contributions

CAMR: Formal Analysis (lead); Methodology (equal); Software (equal); Visualization (equal); Writing – Original Draft Preparation (lead). GRQS: Formal Analysis (supporting); Methodology (supporting). MOCC: Supervision (supporting); ASV: Methodology (supporting). MFC: Methodology (supporting). AMSF: Funding Acquisition (supporting); Investigation (supporting); Methodology (supporting); Resources (supporting); Writing – Review & Editing (supporting). EGV: Formal Analysis (supporting); Methodology (supporting); Software (equal); Visualization (equal). JEMH: Formal Analysis (supporting); Methodology (supporting); Supervision (supporting). MFN: Funding Acquisition (lead); Investigation; Methodology (equal); Project Administration (lead); Resources (lead); Supervision (lead); Writing – Review & Editing (lead).

## Supplementary Material

*(see files at* https://zenodo.org/records/17192305)

**Supplementary Tables S1–S5**. Categorization of sRNA genomic localization across strains. Each table presents the classification of sRNAs according to their genomic annotation (antisense, intragenic, intergenic, 5′ UTR, and 3′ UTR). One table is provided per strain, except for USA500, whose annotation was propagated from USA300, as reported in the original study (Tomlinson, Malof and Shaw, 2021).

**Supplementary Tables S6–S11. Differential gene expression (DEG) between biofilm and planktonic conditions**. Tables list log2 fold changes (log2FC) for each strain, with positive values indicating higher expression in biofilm relative to planktonic growth.

**Supplementary Table S12. Cross-lineage module assignment of genes from co-expression analysis**. For each row, the table lists the locus IDs for Bmb9393 (ST239), N315 (USA100), MRSA252 (USA200), LAC (USA300/USA500), and MW2 (USA400); the

final column reports the WGCNA module (color name) assigned to that locus. Entries include sRNAs (conserved in all strains) and CDSs (blank cells indicate that no CDS homolog was identified in the corresponding strain).

**Supplementary Table S13. Hub genes identified within co-expression modules**. For each strain, hub genes are listed along with their unique IDs, module membership, correlations of modules with biofilm/planktonic states, and, when available, gene product, protein ID, GO ID, and associated GO terms.

**Supplementary Table S14. Gene Ontology (GO) enrichment of co-expression modules**. GO terms associated with individual modules, highlighting biological functions enriched among the genes within each module.

**Supplementary Table S15. High-confidence mRNA–sRNA interaction pairs**. List of candidate regulatory pairs retained after all filtering steps (see Methods). Because of these filters—including cross-tool thresholds, WGCNA co-assignment, and cross-strain expression-consistency—not all antisense (cis) sRNA–CDS pairs were retained; antisense pairs that failed any criterion were excluded and therefore do not appear here.

**Supplementary Table S16.** Metadata of RNA-seq samples used in this study. This table lists the RNA-seq samples from GEO Series GSE163153, including their unique Sample ID (GSM accession) and the corresponding *S. aureus* strains (e.g., N315, MRSA252, USA300_LAC, MW2, NRS385).

| GEO Serie GSE163153 |  |  |  |  |
| --- | --- | --- | --- | --- |
| Sample ID | Strain | Treatment | Time | DOI |
| GSM4973293 | N315 | Biofilm | 24h | 10.1099/mgen.0.000598 |
| GSM4973294 | MRSA252 | Biofilm | 24h | 10.1099/mgen.0.000598 |
| GSM4973295 | USA300_LAC | Biofilm | 24h | 10.1099/mgen.0.000598 |
| GSM4973296 | MW2 | Biofilm | 24h | 10.1099/mgen.0.000598 |
| GSM4973297 | NRS385 | Biofilm | 24h | 10.1099/mgen.0.000598 |
| GSM4973298 | N315 | Planktonic | 24h | 10.1099/mgen.0.000598 |
| GSM4973299 | MRSA252 | Planktonic | 24h | 10.1099/mgen.0.000598 |
| GSM4973300 | USA300_LAC | Planktonic | 24h | 10.1099/mgen.0.000598 |
| GSM4973301 | MW2 | Planktonic | 24h | 10.1099/mgen.0.000598 |
| GSM4973302 | NRS385 | Planktonic | 24h | 10.1099/mgen.0.000598 |

**Supplementary Table S17:**
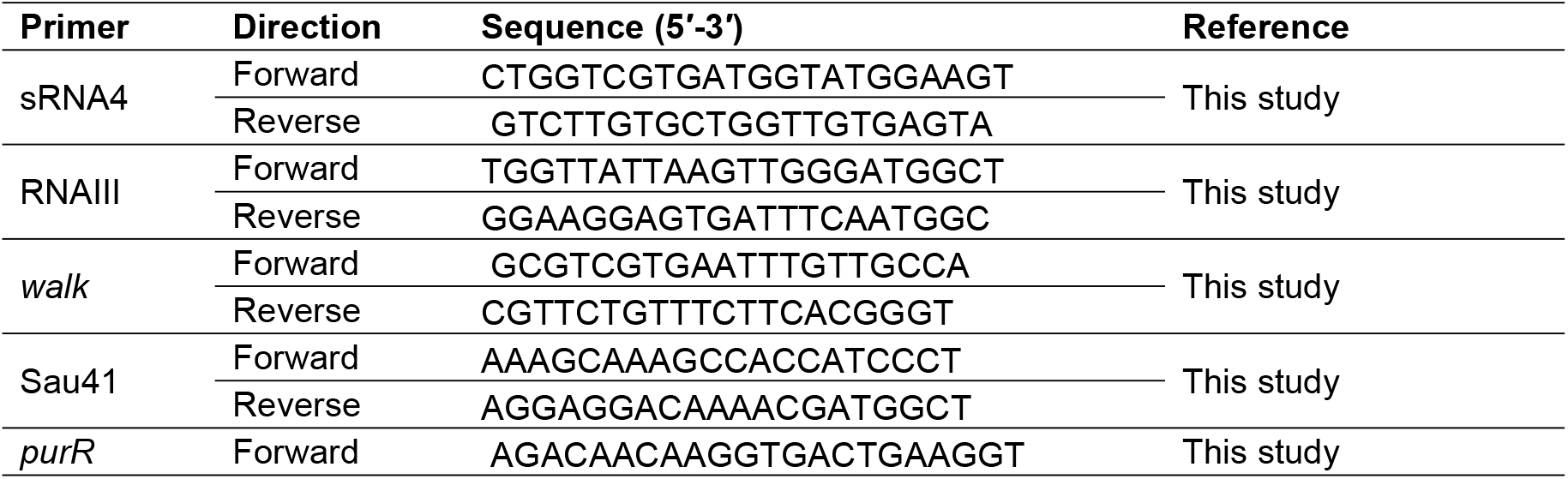
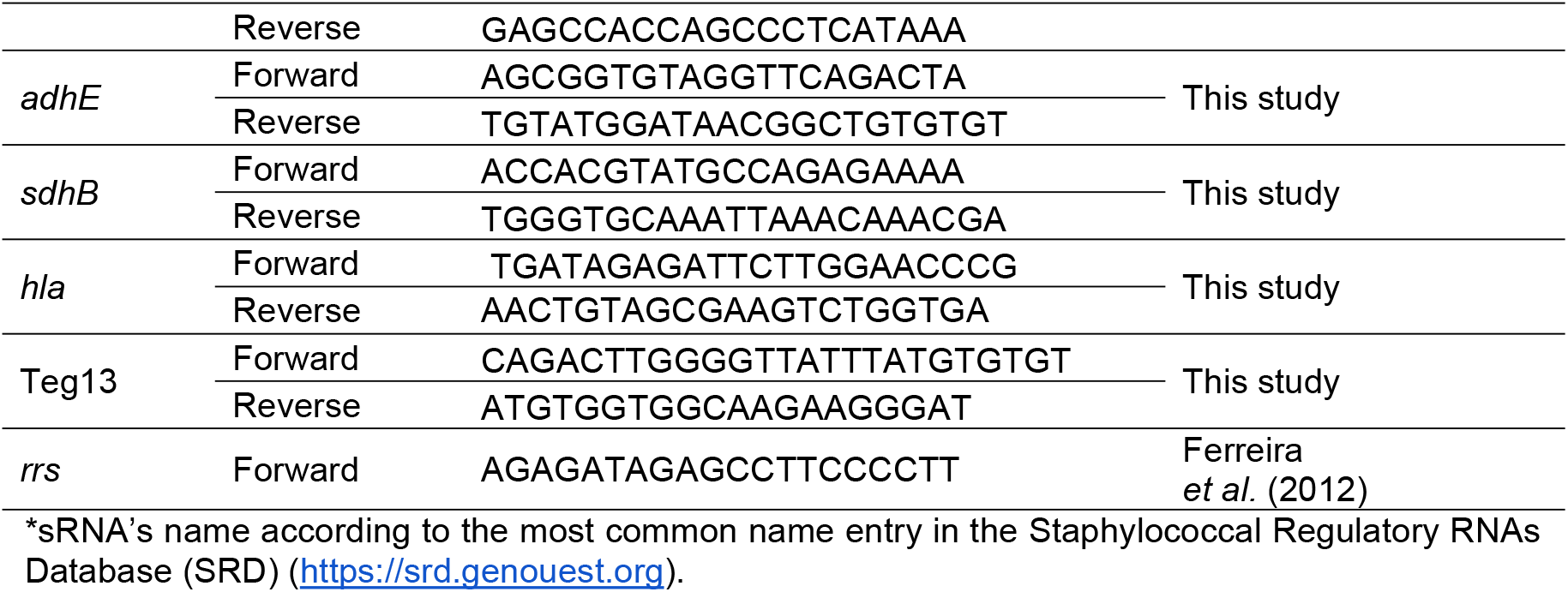
List of RT-qPCR primers.

**Supplementary Table S18.** Predicting Interaction Values for sRNA-mRNA Pairs Reported in the Literature. P-Values from TargetRNA3, INTARna, RNAplex, and Probabilities from sRNARFtarget.

| sRNA | gene | sRNA<br>RF<br>Target | Target<br>RNA<br>3 | intaR<br>NA | RNAp<br>lex | DOI |
| --- | --- | --- | --- | --- | --- | --- |
| artR | *sarT* | 0.48 | 0.21 | 0.1 | 0.6 | DOI: 10.1007/s00430-013-0307-0 |
| rsaA | *mgrA* | 0.6 | 0.16 | 0.35 | 0.47 | DOI: 10.1093/nar/gkx219 |
| rsaA | *ssaA* | 0.58 | 0.02 | 0.07 | 0.47 | DOI: 10.1093/nar/gkx219 |
| rsaA | *flr* | 0.6 | 0.03 | 0.06 | 0.29 | DOI: 10.1093/nar/gkx219 |
| rsaD | *alsS* | 0.53 | 0.13 | 0.04 | 0.4 | DOI: 10.1111/mmi.14418 |
| rsaE | *rocF* | 0.57 | 0.42 | 0.08 | 0.06 | DOI: 10.1093/nar/gky584 |
| rsaE | *opp3*<br>*a/3b* | 0.58 | 0.15 | 0.06 | 0.02 | DOI: 10.1093/nar/gkq462 |
| rsaE | *rocF* | 0.57 | 0.42 | 0.08 | 0.06 | DOI: 10.1093/nar/gky584 |
| rsaE | *rocD* | 0.55 | 0.37 | 0.1 | 0.01 | DOI: 10.1093/nar/gky584 |
| rsaE | *sucD* | 0.55 | 0.04 | 0.14 | 0.04 | DOI: 10.1093/nar/gkp668 |
| rsaE | *sucC* | 0.57 | 0.04 | 0.12 | 0.02 | DOI: 10.1093/nar/gkp668 |
| rsaE | *sa08*<br>*73* | 0.59 | 0.07 | 0.05 | 0.03 | DOI: 10.1093/nar/gky584 |
| rsal | *icaR* | 0.46 | 0.4 | 0.04 | 0.71 | DOI: 10.15252/embj.201899363 |
| RNAII<br>I | *coa* | 0.57 | 0.18 | 0.41* | 0.66 | DOI: 10.1371/journal.ppat.1000809 |
| RNAII<br>I | *ecb* | 0.48 | 0.25 | 0.15 | 0.68 | DOI: 10.1042/BC20070137 |
| RNAII<br>I | *eap* | 0.45* | 0.19 | 0.1 | 0.54 | DOI: 10.1099/mic.0.27902-0 |
| RNAII<br>I | *esxA* | 0.48 | 0.7 | 0.13 | 0.65 | DOI: 10.1038/s41467-022-31173-y |
| RNAII<br>I | *hla* | 0.52 | 0.5 | 0.31 | 0.53 | DOI: 10.1002/j.1460-2075.1995.tb00136.x |
| RNAII<br>I | *ltaS* | 0.53 | 0.09 | 0.19 | 0.62 | DOI: 10.1002/jobm.201400313 |
| RNAII<br>I | *murQ* | 0.53 | 0.58 | 0.36 | 0.56 | DOI: 10.1038/s41467-022-31177-8 |
| RNAII<br>I | *rot* | 0.49 | 0.03 | 0.1 | 0.67 | DOI: 10.1111/j.1365-2958.2006.05292.x |
| RNAII<br>I | *sbi* | 0.51 | 0.67 | 0.1 | 0.64 | DOI: 10.1093/nar/gku119 |
| RNAII<br>I | *spa* | 0.5 | 0.2 | 0.24 | 0.5 | DOI: 10.1038/sj.emboj.7600572 |
| RNAII<br>I | *lytM* | 0.56 | 0.17 | 0.24 | 0.6 | DOI: 10.1002/jobm.201100426 |
| sprD | *sbi* | 0.5 | 0.4 | 0.18 | 0.76* | DOI: 10.1371/journal.ppat.1000927 |
| SprX | *clfB* | 0.5 | 0.04 | 0.16 | 0.27 | DOI: 10.1016/j.micres.2021.126785 |
| SprX | *ecb* | 0.46 | 0.14 | 0.12 | 0.43 | DOI: 10.1160/TH05-05-0306 |
| SprX | *spoVG* | 0.52 | 0.89* | 0.23 | 0.53 | DOI: 10.1038/s41598-021-86376-y |
| SprX | *walR* | 0.51 | 0.1 | - | 0.56 | DOI: 10.1016/j.micres.2021.126785 |
| Teg4<br>1 | *psmα* | 0.52 | 0.44 | 0.28 | 0.42 | DOI: 10.1128/mBio.02484-18 |
| Teg4<br>9 | *sarA* | 0.55 | 0 | 0.13 | 0.49 | DOI: 10.1111/mmi.14919 |

\*sRNA's name according to the most common name entry in the Staphylococcal Regulatory RNAs Database (SRD) (https://srd.genouest.org).

## Supplementary Figures

**Figure S1.**
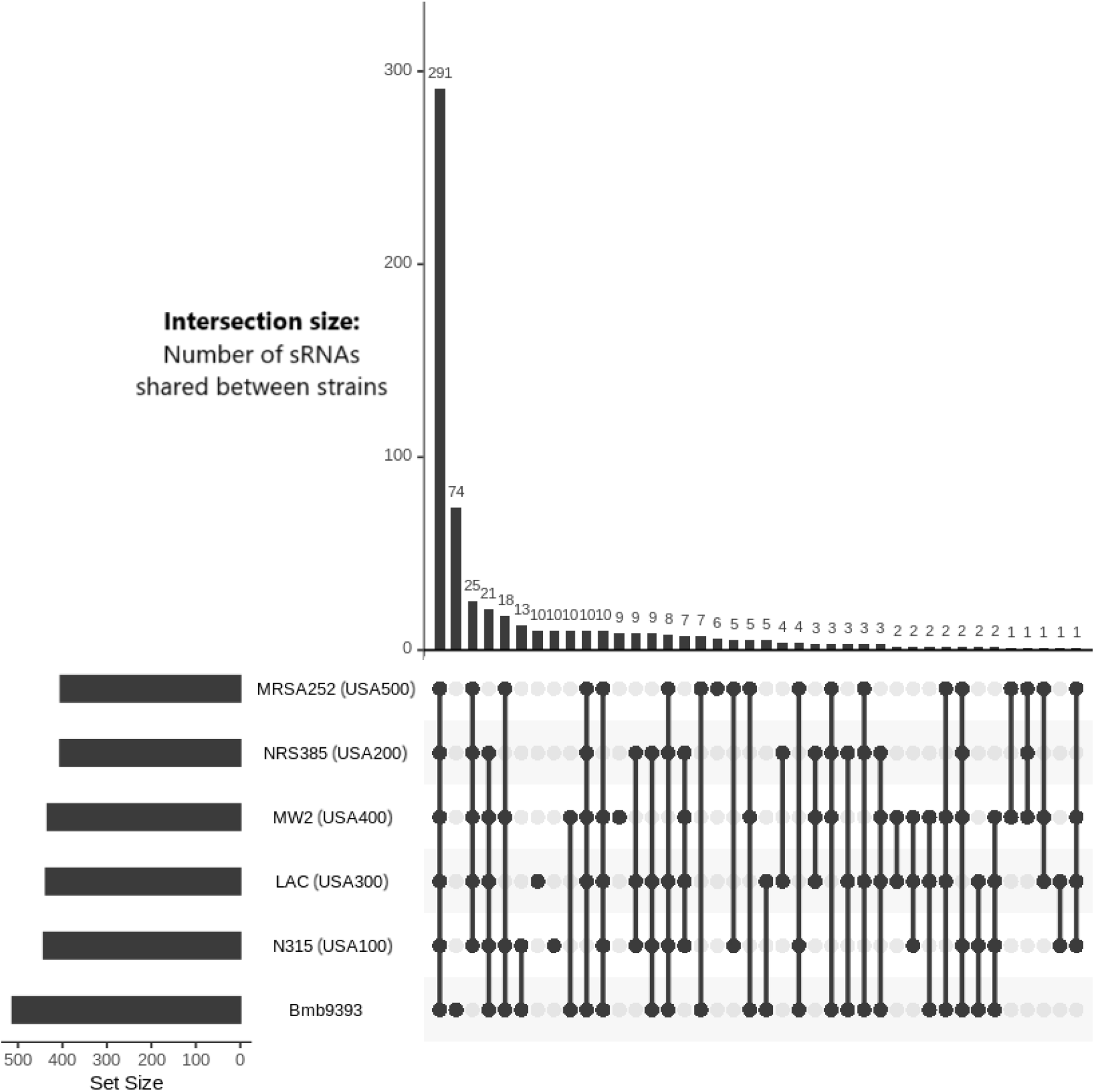
Quantification of the sRNAs with normalized expression levels. (TMM normalized, converted to RPKM) exceeding 10 reads across the studied strains. The horizontal bars on the left represent the total number of sRNAs detected in each strain, while the vertical bars on the top denote the number of sRNAs shared between specific combinations of strains. The connected dots in the matrix below indicate the strain combinations contributing to each intersection set.

**Figure S2.**
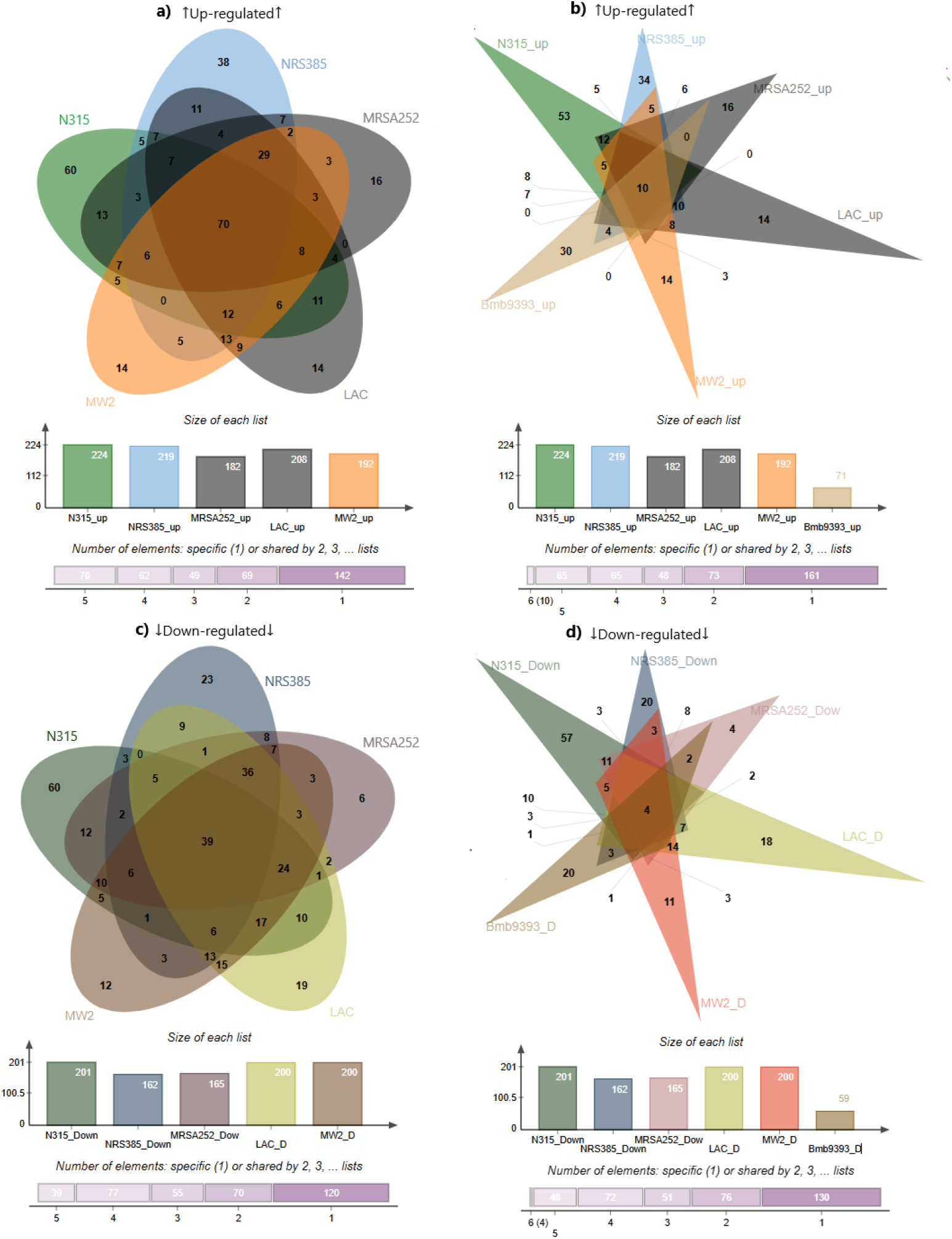
Differentially expressed sRNAs across strains. Venn diagrams summarize sRNAs upregulated in biofilm (top row) and downregulated in biofilm (bottom row), defined relative to planktonic conditions. Left panel: USA-lineage strains (N315, NRS385, MRSA252, LAC, MW2). Right panel: all six strains (USA strains plus Bmb9393). Below, a box plot shows the total number of sRNAs identified per strain. A purple segmented bar depicts prevalence categories for unique sRNAs: present in all strains, in 4/5 strains, in 3/5 strains, in 2/5 strains, or in a single strain.

